# Protection of β2GPI Deficient Mice from Thrombosis Reflects a Defect in PAR3-facilitated Platelet Activation

**DOI:** 10.1101/2023.08.23.554547

**Authors:** Paresh P Kulkarni, Ravi Kumar Alluri, Matthew Godwin, Gabriel L Forbes, Alona Merkulova, Aatira Vijay, Maierdan Palihati, Suman Kundu, Young Jun-Shim, Alvin Schmaier, Michael Holinstat, Scott J. Cameron, Keith R McCrae

**Affiliations:** Department of Cardiovascular and Metabolic Sciences, Cleveland Clinic, Cleveland, OH; Division of Hematology and Oncology, Department of Medicine, Case Western Reserve University, Cleveland, OH; Department of Pharmacology, University of Michigan, Ann Arbor, MI; Heart, Vascular and Thoracic Institute, Cleveland Clinic, Cleveland, OH; Taussig Cancer Institute, Cleveland Clinic, Cleveland, OH

## Abstract

**Background:** Antibodies to β2-glycoprotein I (β2GPI) cause thrombosis in antiphospholipid syndrome, however the role of β2GPI itself in regulation of coagulation pathways *in vivo* is not well understood.

**Methods:** We developed β2GPI-deficient mice *(Apoh^-/-^)* by deleting exon 2 and 3 of *Apoh* using CRISPR/Cas9 and compared the propensity of wild-type (WT) and *Apoh^-/-^*mice to develop thrombosis using rose bengal and FeCl_3_-induced carotid thrombosis, laser-induced cremaster arteriolar injury, and inferior vena cava (IVC) stasis models. We also compared tail bleeding times and assessed platelet activation in WT and *Apoh^-/-^* mice in the absence and presence of exogenous β2GPI.

**Results:** Compared to WT littermates, *Apoh*^-/-^ mice demonstrated a prolonged time to occlusion of the carotid artery after exposure to rose bengal or FeCl_3_, and reduced platelet and fibrin accumulation in cremasteric arterioles after laser injury. Similarly, significantly smaller thrombi were retrieved from the IVC of *Apoh^-/-^*mice 48 hours after IVC occlusion. The activated partial thromboplastin time (aPTT) and prothrombin time, as well as aPTT reagent- and tissue factor-induced thrombin generation times using plasma from *Apoh*^-/-^ and WT mice revealed no differences. However, we observed significant prolongation of tail bleeding in *Apoh^-/-^* mice, and reduced P-selectin expression and binding of fibrinogen to the activated α2bβ3 integrin on platelets from these mice after stimulation with low thrombin concentrations; these changes were reversed by exogenous β2GPI. An antibody to PAR3 blocked thrombin-induced activation of WT, but not *Apoh^-/-^* platelets, as well as the ability of β2GPI to restore the activation response of *Apoh^-/-^* platelets to thrombin. β2GPI deficiency did not affect platelet activation by a PAR4-activator peptide, or ADP.

**Conclusions:** In mice, β2GPI may mediate procoagulant activity by enhancing the ability of PAR3 to present thrombin to PAR4, promoting platelet activation at low thrombin concentrations.

**Key Points:**

- β2GPI deficient mice are protected from experimental arterial, venous, and microvascular thrombosis.
- β2GPI deficient mice display prolonged tail bleeding times and reduced PAR3-facilitated platelet activation by low concentrations of thrombin.

## Introduction

Antiphospholipid syndrome (APS) is an autoimmune disorder characterized by arterial and venous thrombosis and/or recurrent fetal loss in the presence of persistent circulating “antiphospholipid” antibodies (APL).^1–3^ While these antibodies have broad specificity, they primarily react with proteins or protein-phospholipid complexes, with reactivity toward β2GPI having been most thoroughly characterized.^4–6^ β2GPI is a 50 kDa plasma glycoprotein consisting of four typical complement control protein (CCP) domains, followed by an atypical, positively-charged fifth domain that mediates phospholipid binding^7,8^. Anti-β2GPI antibodies reactive with domain 1 of β2GPI appear to have a stronger association with thrombosis than those reactive with other domains^9–15^

*In vitro*, β2GPI has been reported to have both anticoagulant and procoagulant properties. Reduced levels of β2GPI in patients with disseminated intravascular coagulation have been assumed to reflect consumption, and thus it was presumed that β2GPI has a role in regulating thrombin generation.^16^ However, conflicting results on the effects of β2GPI on thrombin generation *in vitro* have been reported by several groups.^17–19^ One group reported that β2GPI bound FXI and blocked its activation by thrombin or FXIIa.^20^. Others have observed that β2GPI either inhibits or has no effect on the activation of protein C^17,21^, or impairs the activity of activated protein C (APC)^22^.

Several *in-vitro* studies also suggest a role for β2GPI in regulation of platelet function. β2GPI has been reported to bind to washed platelets and alter platelet cyclic nucleotide metabolism^23,24^, to inhibit the platelet release reaction in response to ADP^25^, and to impair platelet prothrombinase activity.^26^ β2GPI has also been reported to bind thrombin through exosites 1 and 2, prolong thrombin and ecarin clot times, and impair thrombin-induced aggregation of human platelets by inhibiting cleavage of protease activated receptor 1 (PAR-1).^17^

Finally, β2GPI has been reported to stimulate fibrinolysis through enhancement of t-PA activity^27^, among other mechanisms.^28,29^

These results suggest complex interactions of β2GPI with several coagulation pathways but leave uncertainty concerning the net effect of β2GPI on blood coagulation *in vivo*, where interactions with blood cells, plasma proteins and the vascular wall occur. A recent report suggested that *Apoh-/-* mice displayed a prothrombotic phenotype, suggesting that β2GPI functions as a natural anticoagulant in mice.^30^ However, control mice in these studies were not littermates, and were obtained from a separate vivarium, raising concerns about these conclusions.

A thorough analysis of the role of β2GPI in coagulation *in vivo* is essential to developing a framework to better understand the pathogenesis of APS. To address this issue, we used CRISPR/Cas9 to generate mice deficient in β2GPI, deleting the same exons targeted in previously-reported *Apoh^-/-^* mice.^18^ These mice were viable, with no significant phenotype under non-stressed conditions. However, using four distinct models of thrombosis performed in three different laboratories by blinded observers, we observed that β2GPI deficiency was protective against arterial, venous and microvascular thrombosis. Our studies suggest that these outcomes reflect enhancement of platelet activation at low thrombin concentrations by β2GPI.

## MATERIALS AND METHODS

Materials used in these studies and their sources are listed in **Supplementary Table 1.**

### Generation of Apoh-/- mice

The overall strategy for generation of *Apoh-/-* mice, including guide RNA design, is depicted in **Figure 1A and 1B**. Supplementary Figure 1 demonstrates the experimental validation of guide RNA cleavage sites within their targeted oligonucleotide sequences.

**Figure 1:**
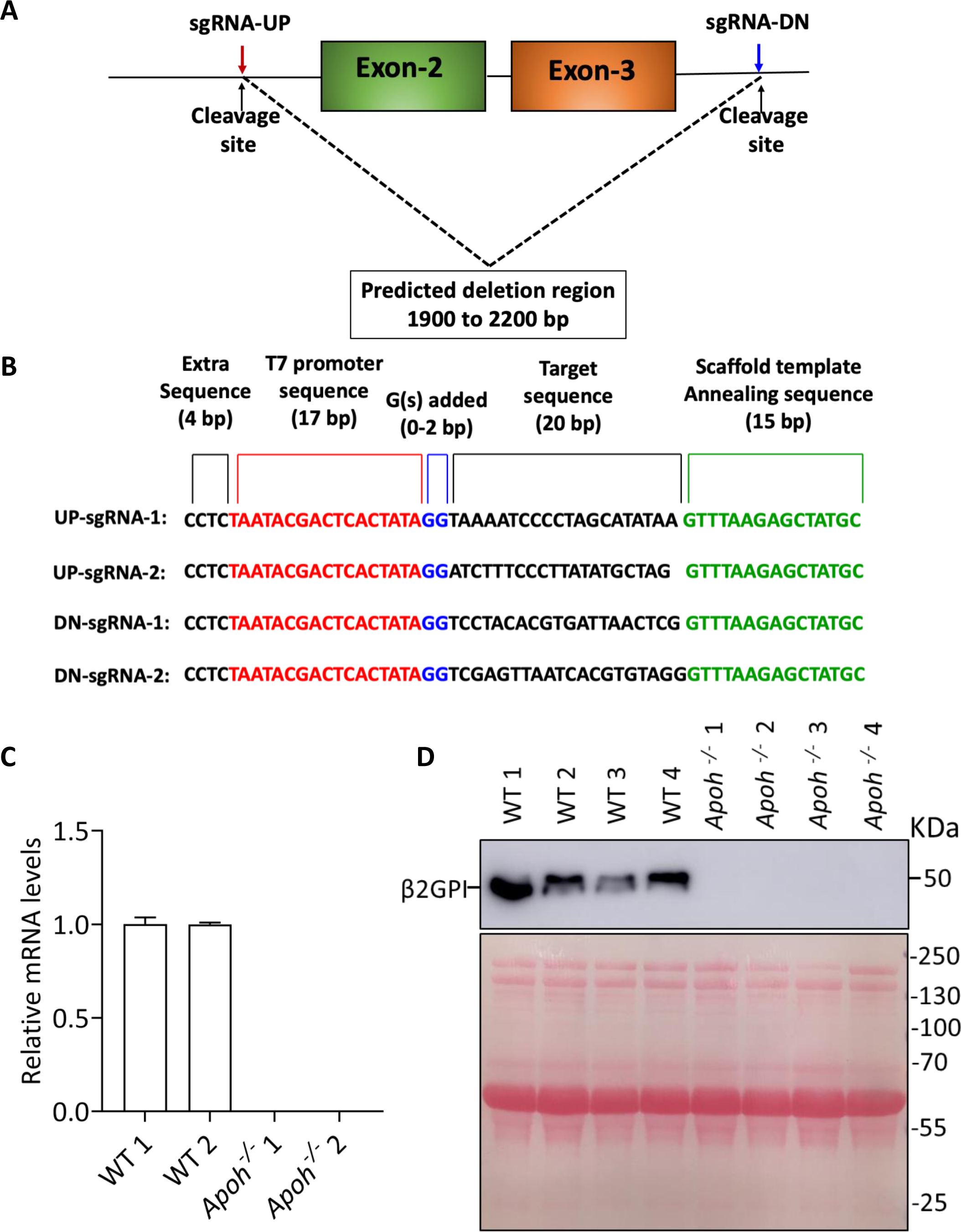
CRISPR/Cas9 mediated *Apoh* deletion in mice. (**A**) The genomic sequence of *Apoh* in C57BL/6J mice was used to generate guide RNAs (sgRNAs) targeting regions upstream of exon 2 and downstream of exon 3. The two best sgRNAs generated for the region upstream of exon-2 (sgRNA-UP-1, sgRNA-UP-2), and downstream of exon 3 (sgRNA-DOWN-1, sgRNA-DOWN-2) were selected. (**B**) Templates for *in vitro* transcription of sgRNAs were designed containing an extra four 5’ base pairs (black) followed by a T7 promoter (red), target sequence (black) and a 15 base pair scaffold sequence (green). **(C)** *Apoh* mRNA levels in liver from WT and *Apoh^-/-^* mice**. (D)** Immunoblot (upper panel) for β2GPI in WT and *Apoh^-/-^* mice. Lower panel = total protein stain with Ponceau S.

### Characterization of Apoh^-/-^ mice

*Apoh^-/-^* mice were viable with no obvious phenotypic abnormalities. The frequency of *Apoh^-/-^*, *Apoh^+/-^*and wild-type (WT) offspring followed expected Mendelian frequencies.

The genotyping strategy for *Apoh^-/-^* mice is depicted in **Supplementary Figure 2A** and genotyping results from representative mice in Supplementary Figure 2B. Since the liver is the primary site of *Apoh* expression, we also confirmed the absence of *Apoh* mRNA in liver of *Apoh^-/-^* mice.

To assess β2GPI in plasma, five microliters of plasma from *Apoh^-/-^* and *Apoh^+/+^* (WT) mice was diluted 1:10 in Laemmli sample buffer containing *β*-mercaptoethanol and loaded onto a 10% SDS-PAGE gel. Proteins were transferred to PVDF, and immunoblotted using an affinity-purified, polyclonal rabbit anti-β2GPI antibody. Bound antibody was detected using peroxidase-conjugated goat-anti rabbit-IgG followed by detection using SuperSignal West Pico Plus chemiluminescent substrate.

### Coagulation assays

Blood for coagulation assays (0.35 ml) was drawn from the IVC using a 1 ml tuberculin syringe containing 0.035 ml of 3.2% sodium citrate and immediately transferred into 1.5 ml polypropylene microcentrifuge tubes. Tubes were centrifuged for 2 min in an Eppendorf microcentrifuge and the plasma stored at -80° C.

The activated partial thromboplastin time (aPTT) was performed as previously described.^31^ Briefly, 50 µl of citrated plasma was mixed with 50 µl of pre-warmed kaolin-based aPTT reagent in a glass cuvette, and incubated in a 37° C water bath for 5 min. Clotting was initiated by adding 50 µl of 35.3 mM CaCl_2_ and the clot formation endpoint was determined by visual observation while continuously rocking the cuvette. The prothrombin time (PT) was performed by adding 100 µl of pre-warmed PT reagent to 50 µl citrated plasma in a glass cuvette and the clotting time determined as described for the aPTT assay.^31^

### Thrombin generation times

Contact activation and tissue factor-induced thrombin generation times (TGT) were performed in Nunclon Delta Surface 96 well microtiter plates (Thermo Scientific).^31^ The assay was performed in 25 mM HEPES, 150 mM NaCl pH 7.4 containing 2 mg/ml bovine serum albumin (BSA) in a total volume of 100 μl. The contact activation TGT was performed as follows: In a well containing 45 μl of buffer, 15 μl of mouse plasma was added followed by 10 μl of phospholipids (5 μM final concentration), 10 μl of Z-Gly-Gly-Arg-AMC (0.42 mM final concentration), and 10 μl of aPTT reagent. The assay was initiated by the addition of 10 μl of 30 mM CaCl_2_ followed by insertion of the microplate into a Molecular Devices SpectraMaxi3x fluorometer at 23^°^C. The tissue factor TGT was performed similarly: to wells containing 35 μl of buffer, 30 μl of mouse plasma was added, followed by 10 μl of phospholipids (5 μM final concentration), 10 μl of Z-Gly-Gly-Arg-AMC (0.42 mM final concentration), and 5 μl of a 1:10 dilution of Innovin stock (∼0.25 pM). The assay was initiated by the addition of 10 μl of 30 mM CaCl_2_, and reactions monitored using a Molecular Devices SpectraMaxi3x fluorometer at 23^°^C. Each tissue factor TGT included a companion sample performed without addition of Innovin. Fluorescent readings were performed at an excitation of 360 nm and an emission of 440 nm for 90 min. Generated data were analyzed in PRISM (GraphPad Software LLC, San Diego, CA). In tissue factor TGT assays, the substrate hydrolysis of the companion sample was first subtracted from the tissue factor sample before data calculation to remove any contribution of contact activation during sample incubation. Initial rates of thrombin generation were demonstrated for each contact activator or tissue factor induced TGT sample and first derivative TGT curves and area under the curve (AUC) were generated using PRISM 9.3.1. Each TGT curve and AUC is the mean + SEM of 5 independent experiments using plasma from 5 different mice.

### In vivo thrombosis studies

Thrombosis in *Apoh*^-/-^ mice and WT littermates was compared using four different assays, two assessing arterial, one arteriolar, and one venous thrombosis. For the rose bengal-induced carotid artery thrombosis assay, 8- to 12-week-old mice were anesthetized with sodium pentobarbital and placed under a dissecting microscope (Nikon SMZ-2T; Mager Scientific, Inc., Dexter, MI) in the supine position. The right common carotid artery was exposed by a midline surgical incision and a Doppler flow probe (Model 0.5 VB; Transonic Systems, Ithaca, NY) was placed under the vessel to measure blood flow using a flowmeter (model T106, Transonic Systems, Ithaca, NY). One hundred twenty microliters of rose bengal (4,5,6,7-Tetrachloro-3’,6’-dihydroxy-2’,4’,5’,7’-tetraiodo-spiro[isobenzofuran-1(3H),9’ [9H]xanthen]-3-one disodium salt) at a concentration of 50 mg/kg in 0.9% NaCl was injected into the tail vein^32,33^. A green laser light (Melles Griot, Carlsbad, CA) was applied 6 mm from the carotid artery at a wavelength of 540 nm to generate reactive oxygen species from photoactivation of rose bengal. Blood flow was continuously monitored from the initiation of photochemical injury, and the time to complete occlusion was determined based on the time to onset of stable cessation of blood flow maintained for 20 minutes. Data was acquired and analyzed using the Windaq software program (Windaq; DATAQ Instruments, Akron, OH).

We also assessed FeCl_3_-induced carotid artery thrombosis. Briefly, mice (8-12 wk) were anesthetized as for rose bengal studies, and the common carotid artery exposed. An ultrasonic miniature Doppler flow probe (Model 0.5 VB; Transonic Systems, Ithaca, NY) was placed under the vessel, and a 1 x 1 mm piece of Whatman filter paper soaked in 4% FeCl_3_ was applied to the carotid artery for one minute. The filter paper was then removed, the vessel was washed with saline, and blood flow was monitored using a flowmeter (Transonic model T106, Transonic Systems). Data was acquired and analyzed as for rose bengal assays.

Microvascular arteriolar thrombosis was assessed as previously described.^34^ Briefly, *Apoh^-/-^* male mice or their WT littermates (12-14 weeks of age) were anesthetized by intraperitoneal injection of ketamine/xylazine (100 and 10 mg/kg respectively). A jugular vein catheter was established to inject antibodies (DyLight 488 anti-GPIbβ, Emfret Analytics, Eibelstadt, Germany and anti-mouse fibrin antibody, a kind gift from Dr R. Camire, Children’s Hospital Of Philadelphia, labeled with Alexa Fluor 647). The cremaster muscle was surgically prepared under a dissecting microscope and superfused throughout the experiment with preheated bicarbonate saline buffer (132 mM NaCl, 4.7 mM KCl, 1.2 mM MgS0_4_, 2 mM CaCl_2_, 18 mM NaHCO_3_). The cremaster muscle arteriole (30-50 µm diameter) was exposed to high intensity 532 nM laser pulses (Ablate! Photo-ablation system; Intelligent Imaging Innovations, Denver, CO) to injure the arterial wall and induce thrombus formation. Multiple laser injuries were performed in each mouse, with each injury upstream of all prior injuries. Images were acquired in real-time at 0.2s intervals using a 63x water-immersion objective, with a Zeiss Axio Examiner Z1 multichannel fluorescent microscope equipped with a solid laser launch system and high-speed sCMOS camera. The dynamic process of platelet accumulation and fibrin formation was analyzed by changes in the mean fluorescence intensity over time after subtracting fluorescent background defined on an uninjured section of the vessel using SlideBook 6.3 software (Intelligent Imaging Innovations).

We also compared venous thrombosis in WT and *Apoh-/-* mice following occlusion of the IVC^35^. Eight- to 12-week-old mice were anesthetized and placed on a heated pad at 37°C. The abdomen was incised, and the intestine exteriorized to allow visualization of the IVC. IVC side branches were ligated with 7−0 prolene sutures while back branches from the renal veins to the iliac bifurcation were cauterized. The IVC was then separated from the aorta and ligated using a 7−0 prolene suture. The abdominal cavity was closed using 5-0 vicryl sutures and adhesive skin glue. Mice were sacrificed 48 hours later and IVC thrombi were harvested, weighed, and analyzed.

### Platelet activation studies

Retro-orbital blood was obtained from 8-12-week-old mice using a capillary tube and immediately placed into (1:9) acid citrate dextrose (ACD) and then diluted (1:1) with modified Tyrode’s buffer. Samples were centrifuged for 15 minutes at 100 g, and the supernatant centrifuged again at the same speed for 10 minutes. The supernatant from the second centrifugation was collected into a fresh tube, supplemented with 3 μM prostaglandin E_1_ (PGE_1_) and centrifuged at 800 g for 7 minutes. The washed platelet pellet from this centrifugation was resuspended in fresh modified Tyrode’s buffer.

Platelets were stimulated with thrombin (0.1, 0.25 or 0.5 U/ml), AYPGKF (PAR4 activating peptide; 100, 250, 500 µM) or ADP (2, 5 or 10 µM) for 15 minutes and analyzed for activation markers by flow cytometry. To confirm the role of β2GPI in platelet activation, *Apoh*^-/-^ platelets were supplemented with plasma-derived β2GPI (40 to 200 µg/ml) before agonist stimulation. In experiments carried out to evaluate the role of PAR3-facilitated signaling in thrombin-induced platelet activation, WT and *Apoh*^-/-^ platelets with or without β2GPI supplementation were pre-incubated for 15 minutes with an antibody reactive with amino acid sequence 31-47 of PAR3 (clone 8E8; 10 µg/ml), which targets the thrombin-binding site. In another experiment to rule out the contribution of glycoprotein Ib (GPIb) signaling in thrombin-induced platelet activation, WT and *Apoh*^-/-^ platelets with or without β2GPI supplementation were pre-incubated for 30 minutes with either O-sialoglycoprotein endopeptidase (OSGE; 80 µg/ml) that proteolytically removes the extracellular domain of GPIbα from the platelet surface^36^ or fibrinogen-γ-peptide (200 µM) that prevents thrombin-GPIbα interaction by blocking thrombin exosite II^17^. After stimulation, 1 µL of anti-CD62P-FITC (P-selectin antibody) or Alexa Fluor 488-conjugated fibrinogen were incubated with 100 µL platelet suspension in separate tubes for 30 minutes in the dark. Platelets were then fixed by the addition of an equal volume of 2% formalin, and quantification of platelet surface P-selectin and fibrinogen binding was performed by acquiring and analyzing 10,000 platelet events from each sample on a flow cytometer (Accuri Flow Cytometer, BD Biosciences)^37,38^. FlowJo (FlowJo LLC, Ashland, Oregon) software was used to process and analyze flow cytometry data.

### Tail bleeding

Mouse tail clip bleeding assays were performed as previously described^39^. Briefly, 8-12-week-old mice were anesthetized, and the tail transected approximately 3 mm from the tip. The tail was then placed in a 50 ml falcon tube containing saline pre-warmed to 37°C. The time to cessation of bleeding was determined visually. Red cells collected in the saline solution were pelleted by centrifugation and lysed prior to colorimetric determination of hemoglobin content using Drabkin’s Reagent.

### Measurement of protein C activity

Protein C activity in WT and *Apoh*^-/-^ plasma was performed using the Chromogenix Coamatic Protein C activity kit (Diapharma). Briefly, 50 μl of protein C activator (Agkistrodin Contortrix Contortrix venom) was added to 25 μl of plasma diluted 1:3 in deionized H_2_O, and incubated at 37° C for 10 min. The protein C substrate, L-Pyroglutamyl-L-prolyl-L-arginine p-Nitroaniline hydrochloride (S-2366) was added and the sample incubated for an additional 10 min at 37° C. Reactions were terminated by addition of 20% acetic acid, and the absorbance at 405 nM was recorded. Protein C activity was also calibrated using a standard curve generated using a normal human plasma calibrator.

### Statistical analyses

Fold change in relative mRNA expression levels were calculated using the comparative Ct method.^40,41^ All coagulation assays were performed twice with data points derived from triplicate samples; data are presented as mean ± standard deviation (SD) unless otherwise indicated. Statistical differences between groups were determined using the student’s t-test for unpaired samples. Statistical differences of 3 or more groups were compared by ANOVA. For thrombin generation times, the change in slope as a function of time, and area under the curve (AUC) was determined using GraphPad PRISM version 9.3.1. P values < 0.05 were considered statistically significant.

## RESULTS

### Generation and characterization of *Apoh*^-/-^ mice

A clonal population of mice derived from offspring of lineage 7 (**Supplementary Table 4**), with a 1937 nucleotide deletion encompassing exons 2 and 3 of *Apoh*, was selected for these studies. To confirm the absence of *Apoh* in these mice, we observed that *Apoh* mRNA was not detectable in liver, normally the source of β2GPI production (**Figure 1C**). We also probed plasma for β2GPI, finding it to be present in WT but absent in *Apoh^-/-^* mice (**Figure 1D**).

To determine whether *Apoh* deficiency had any effect on routine blood counts, hemograms were performed on a cohort of *Apoh^-/-^* mice and their WT littermates. There were no significant differences observed in levels of hemoglobin, platelet or leukocyte counts, or leukocyte differential counts (**Supplementary Table 5**).

### *Apoh^-/-^* mice are protected from arterial and venous thrombosis

We used four different assays, performed in a blinded manner in three different laboratories, to compare the thrombotic propensity of WT and *Apoh^-/-^* mice. In initial studies, we found that the time to carotid artery occlusion, defined by cessation of blood flow, was significantly prolonged in *Apoh^-/-^*mice (36.6 ± 6.31 min) compared to WT littermates (18.9 ± 3.69 min) after photochemical injury induced by rose bengal (P <0.0001). (**Figure 2A**). Prolonged occlusion times were observed in both male and female *Apoh*^-/-^ mice compared to WT littermates of the same gender (**Figure 2B**).

**Figure 2:**
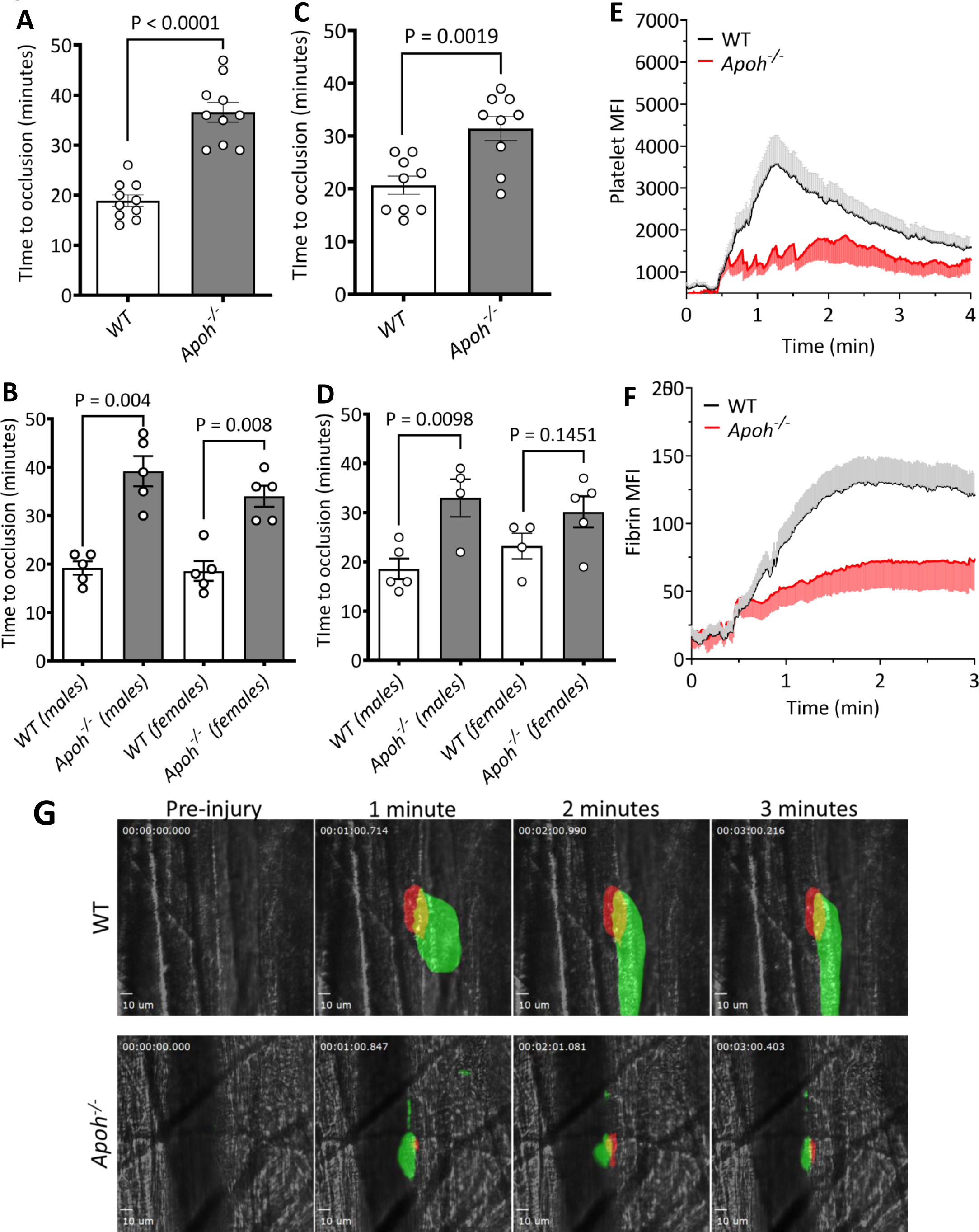
*Apoh^-/-^* mice are protected from arterial and microvascular thrombosis. (A) Time to carotid artery occlusion in WT and *Apoh^-/-^* mice following photoactivation of rose bengal. (B) Data from (A) by gender. (C) Time to carotid artery occlusion in WT and *Apoh^-/-^* mice following exposure to 4% FeCl_3_. (D) Data from (C) by gender. (E and F) Thrombus growth in real time after laser-induced injury of cremaster muscle arterioles of WT or *Apoh^-/-^* mice (n = 4-10 per group) by measuring the mean fluorescence intensity (MFI) of platelet (E) and fibrin (F) acc umulation. (G) Representative images of platelet (anti-GPIb; green) and fibrin (red) accumulation at the indicated times after laser-induced injury of cremaster muscle arterioles in WT or *Apoh^−/−^* mice acquired using a 3I intravital microscopy system on a Zeiss Examiner at ×63 magnification.

We also examined carotid occlusion times in *Apoh^-/-^*and WT littermates using a FeCl_3_-induced arterial thrombosis model. As observed with rose bengal, *Apoh^-/-^* mice displayed a significantly prolonged time to occlusion (31.44 ± 6.98 min) compared to WT littermates (20.67 ± 5.19 min) (P = 0.0019) (**Figure 2C**). Male *Apoh^-/-^* mice also displayed a significantly longer time to occlusion than WT littermates in the FeCl_3_ assay (P = 0.0098); a similar trend was observed in female mice (time to occlusion in *Apoh^-/-^*females = 30.20 ± 7.05 minutes compared to 23.25 minutes in WT females), though these differences did not achieve statistical significance (P = 0.1451) due to the small sample size (**Figure 2D**).

In order to assess the early events occurring during arteriolar thrombosis, *Apoh^-/-^* and WT littermates were subjected to intravital imaging of laser injury-induced thrombosis in cremasteric arterioles. We found that both platelet accumulation (**Figure 2E and G**) and fibrin formation (**Figure 2F and G**) were significantly impaired in *Apoh^-/-^*mice (**Supplementary video 1**) compared to WT littermates (**Supplementary video 2**). Examination of platelet adhesion during early stages of thrombus formation demonstrated a “sawtooth” pattern of platelet adhesion kinetics to the injured arteriolar wall in *Apoh^-/-^* mice (**Figure 2E**), resulting from formation of an unstable platelet plug that separated from the vascular wall soon after formation. This pattern might result from failure of sufficient platelet activation to support thrombus expansion during the early stages of thrombus formation.

To determine whether the absence of β2GPI has similar effects on venous thrombosis, we assessed thrombi developing after occlusion of the IVC. After 48 hours of IVC ligation, thrombi that developed in *Apoh^-/-^* mice were significantly smaller than those in WT littermates [16.94 ± 7.13 mg (n = 16) versus 27.69 ± 6.68 mg (n = 15), respectively (P = 0.0002)] (**Figure 3A and B**). When analyzed by gender, both *Apoh^-/-^*males and *Apoh^-/-^* females had significantly smaller thrombi than corresponding WT littermates of the same gender (*Apoh^-/-^* males = 18.43 ± 8.43 mg, WT males = 29.45 ± 6.31 mg, P = 0.0022; *Apoh^-/-^* females = 14.45 ± 3.52 mg, WT females = 20.63 ± 0.30 mg P = 0.0221) (**Figure 3C**).

**Figure 3:**
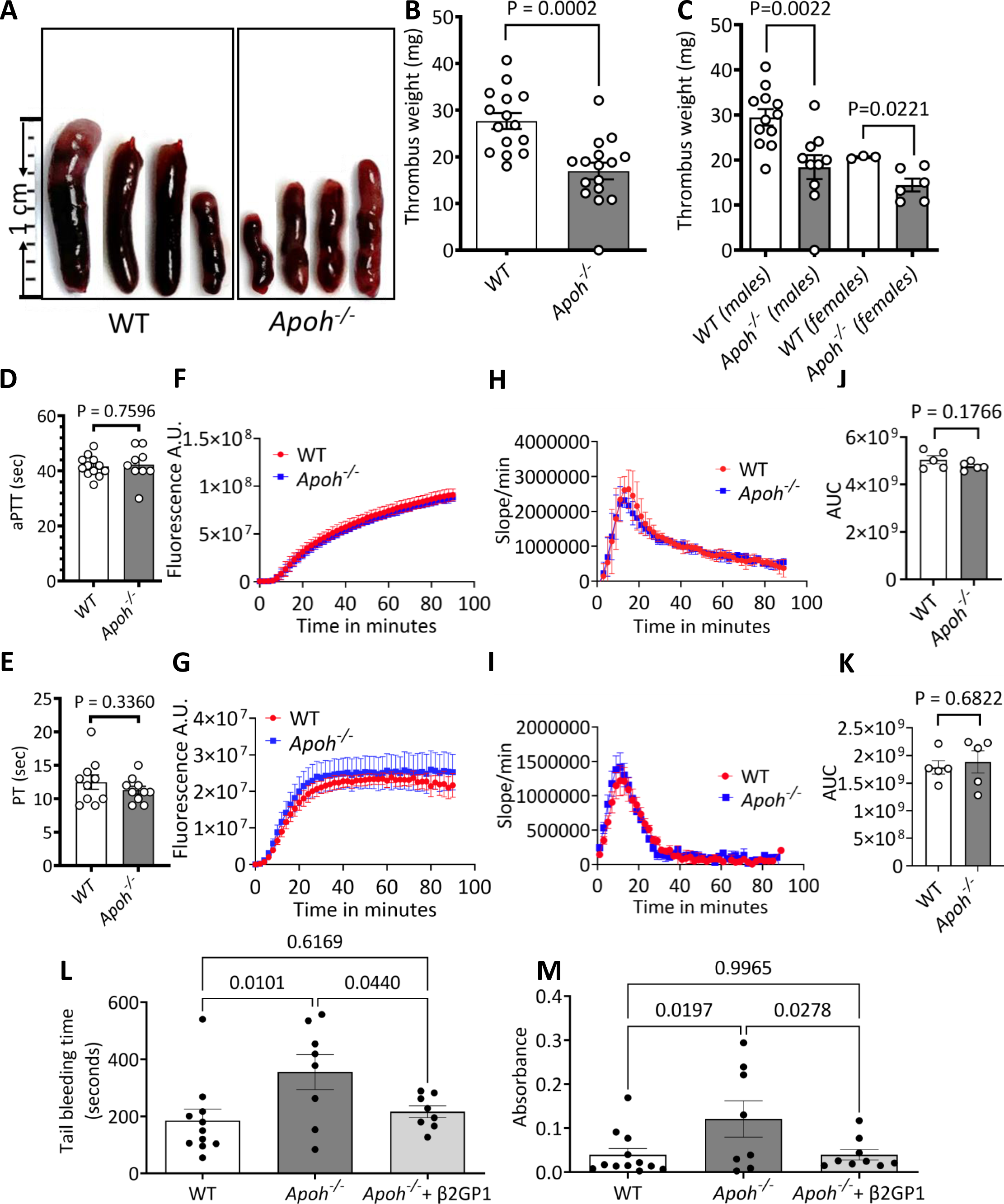
Venous stasis, thrombin generation and tail bleeding. IVC ligation was performed in WT mice or *Apoh^-/-^* littermates. Thrombi were harvested 48 hours after IVC occlusion. (A) Representative thrombi from the IVC of WT and *Apoh^-/-^* mice. (B) Weight of thrombi from WT (n = 15) and *Apoh^-/-^* (n = 16) mice. (C) Data from (B) by gender: WT males (n = 12), WT females (n = 3), *Apoh^-/-^* males (n = 10), *Apoh^-/-^* females (n = 6). (D) Activated partial thromboplastin times. (E) Prothrombin times (F) aPTT reagent-induced thrombin generation. (G) Tissue factor-induced thrombin generation. (H) aPTT reagent-induced thrombin generation rates. (I) TF-induced thrombin generation rates. (J) AUC from (H). (K) AUC from (I). (L) Tail bleeding times in WT (n = 11), *Apoh^-/-^* (n = 8) and *Apoh^-/-^* mice treated with β2GPI (20 mg/kg) (n = 8). (M) Tail bleeding in WT (n = 12), *Apoh^-/-^* (n = 8) and *Apoh^-/-^*mice treated with purified β2GPI (20 mg/kg) (n = 9) quantified by Drabkin’s assay.

### Hemostatic parameters in Apoh^-/-^ mice

To assess the hemostatic mechanisms underlying the hypocoagulability of *Apoh^-/-^* mice, we compared the results of PT and aPTT assays using WT and *Apoh^-/-^* plasma. The mean aPTT was 41.67 ± 3.75 seconds for WT and 42.33 ± 6.08 seconds for *Apoh^-/-^* mice (P = 0.7596) (**Figure 3D**). The mean PT for WT mice was 12.50 ± 3.37 seconds compared to 11.30 ± 1.82 seconds for *Apoh^-/-^* mice (P = 0.3360) (**Figure 3E**).

We also examined thrombin generation times using aPTT reagent and tissue factor to activate the intrinsic and extrinsic pathways of coagulation, respectively. There were no significant differences in either contact activation or tissue factor-induced TGT in plasma from *Apoh^-/-^* and WT mice. Thrombin generation in response to aPTT reagent and tissue factor are depicted in **Figures 3F and 3G**, respectively, and thrombin generation rates in **Figures 3H and 3I**. The AUC determined by these curves is shown in **Figures 3J and 3K**, respectively, with no significant differences observed between WT and *Apoh^-/-^*mice.

Inhibition of protein C activity by APL in the presence of β2GPI has been reported as a potential mechanism for thrombosis in APS, though there is comparatively little information on the effect of β2GPI alone on protein C. Therefore, we determined whether altered protein C activity might contribute to the alterations in thrombosis observed in *Apoh^-/-^* mice. (**Supplementary Figure 3A**). When compared to a standard curve prepared with a human plasma calibrator, the percent protein C activity between the two mouse strains was not significantly different (WT, 8.04 ± 3.20; *Apoh^-/-^*, 6.44 ± 2.04) (P = 0.3743) (**Supplementary Figure 3B**).

### Platelet function in *Apoh^-/-^* mice

Since coagulation assays revealed no significant differences between WT and *Apoh^-/-^* mice, we determined whether any differences existed in platelet function between the two strains. Tail bleeding studies revealed significantly prolonged tail bleeding time in *Apoh^-/-^* mice compared to WT littermates, which was corrected by pre-administration of purified β2GPI (20 mg/kg) (**Figure 3L**). Collection of blood from the time of tail cut to the cessation of bleeding also revealed a higher concentration of hemoglobin in the collection from *Apoh^-/-^* mice, consistent with greater blood loss (**Figure 3M**). This was also reversed by pretreatment of mice with β2GPI.

Next, we assessed the response of washed platelets to varying concentrations of thrombin (0.1-0.5 U/ml), using the expression of P-selectin on the platelet surface, as well as the binding of labeled fibrinogen to platelets as markers of platelet α-granule release and α_IIb_β_3_ integrin activation, respectively.^37,38,42^ No significant differences in expression of P-selectin or binding of fibrinogen to resting platelets from WT or *Apoh^-/-^*mice were observed. However, upon activation by low thrombin concentrations, platelets from *Apoh^-/-^* mice expressed significantly less P-selectin and bound significantly less fibrinogen than platelets from WT littermates (**Figure 4A-D**).

**Figure 4:**
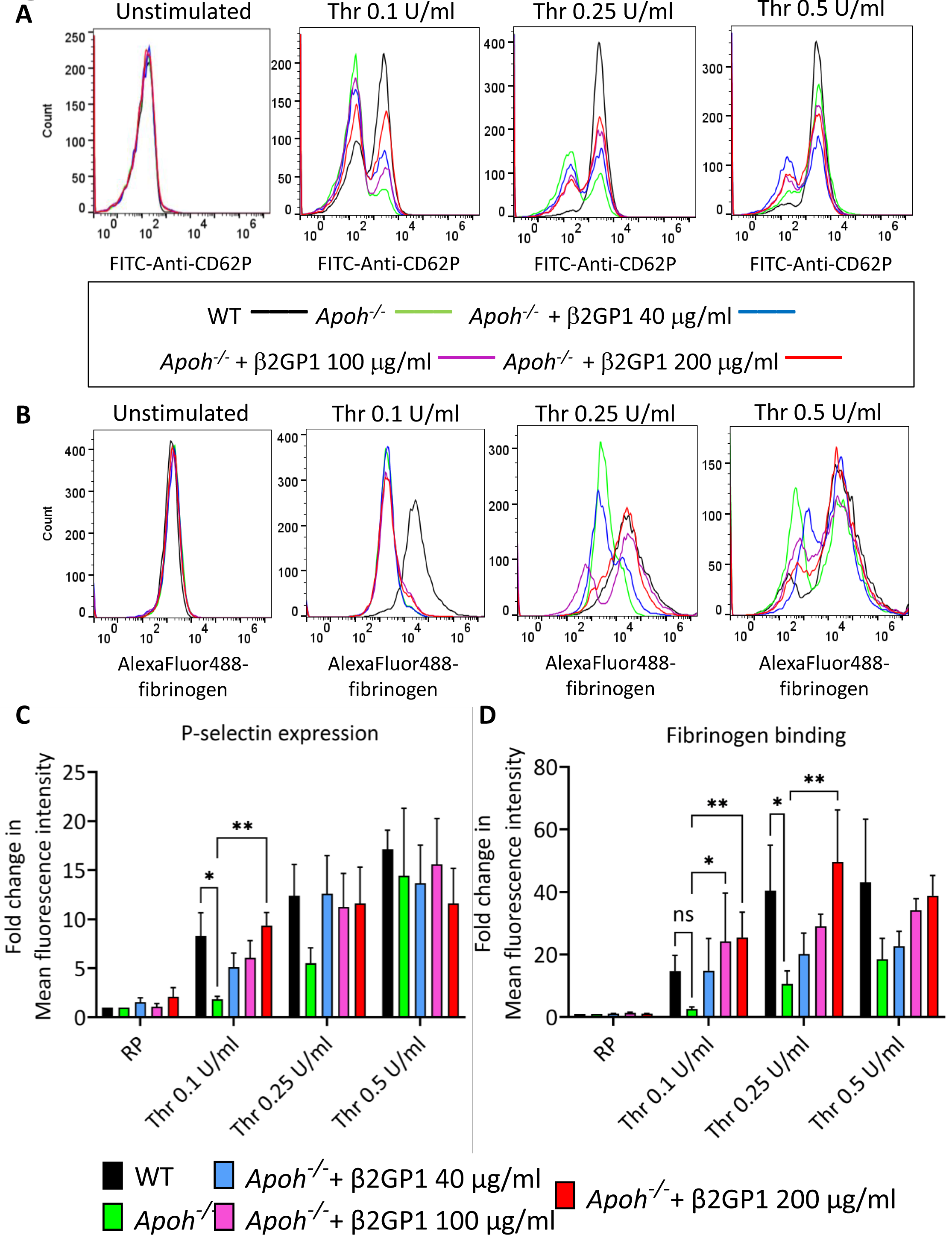
Thrombin-induced activation of WT and *Apoh^-/-^* platelets and the effect of β2GPI. Histogram overlays showing thrombin-induced (A) P-selectin expression and (B) fibrinogen binding of WT platelets in comparison to *Apoh^-/-^* platelets supplemented with β2GPI (40 to 200 µg/ml). (C) and (D) Corresponding bar charts for mean P-selectin expression and fibrinogen binding, respectively.

We reaffirmed that the platelet activation defect in *Apoh^-/-^* mice was a result of β2GPI deficiency by repeating platelet function analyses after supplementing *Apoh^-/-^*platelet suspensions with increasing concentrations of plasma β2GPI (40-200 µg/ml). The defect in low-dose thrombin-induced P-selectin expression (**Figure 4A and C**) and fibrinogen binding (**Figure 4B and D**) of *Apoh^-/-^* platelets compared to WT platelets was reversed by β2GPI. There was a linear dose-response relationship between β2GPI concentration and thrombin-induced activation of *Apoh^-/-^* platelets as assessed by P-selectin expression (**Figure 5A**) and fibrinogen binding (**Figure 5B**).

**Figure 5:**
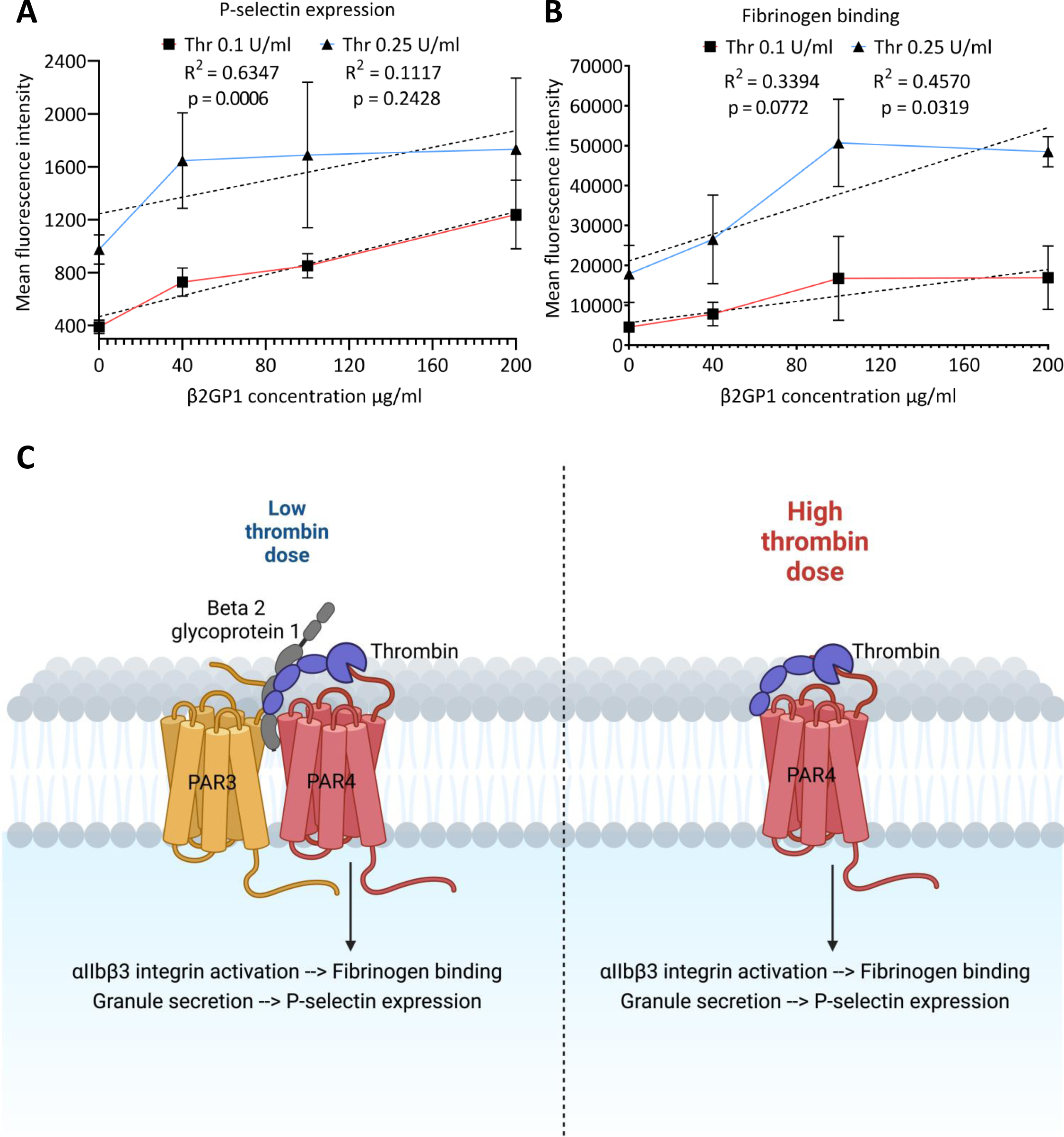
Relationship of β2GPI concentration to (A) P-selectin expression and (B) fibrinogen binding by *Apoh^-/-^*platelets stimulated by low thrombin concentrations. (C) β2GPI may contribute to PAR3-dependent enhancement of PAR4 activation at low thrombin concentrations. In this figure, PAR3 (orange) is shown adjacent to PAR4 (red). Thrombin (blue) binds to the high affinity hirudin-like region of PAR3. β2GPI (gray) may potentially bind to or interact with one or more components of this complex to promote PAR3-PAR4-thrombin interactions and enhance thrombin cleavage of PAR4. At high thrombin concentrations, murine platelets activate through PAR4 in a PAR3 and β2GPI independent manner.

Activation of murine platelets by thrombin is mediated by signaling through protease activated receptor 4 (PAR4), which at low thrombin concentrations is dependent on facilitation of thrombin-PAR4 interaction by PAR3. As *Apoh^-/-^* platelets demonstrated impaired activation only in response to low concentrations of thrombin, we hypothesized that β2GPI is involved in PAR3-facilitated thrombin signaling through PAR4, perhaps through binding thrombin and orienting it to bridge PAR3 and PAR4 more effectively (**Figure 5C**). To test this hypothesis, we adopted a three-pronged strategy to assess the effect of β2GPI deficiency on platelet activation independent of PAR3.

First, we measured activation of WT and *Apoh^-/-^* platelets in the presence or absence of an antibody directed to the thrombin binding site on PAR3. Consistent with our hypothesis and existing dogma, this antibody impaired low-dose thrombin-induced P-selectin expression (**Figure 6A and C**) and fibrinogen binding (**Figure 6B and D**) by WT, but not *Apoh^-/-^* platelets, suggesting that β2GPI is essential for the ability of PAR3 to enhance platelet activation at low thrombin concentrations. Consistent with this observation, there was no significant difference in activation of WT and *Apoh^-/-^*platelets by a low concentration of thrombin in the presence of the PAR3 antibody (**Figure 6**). Similarly, we observed that the ability of β2GPI to enhance P-selectin expression and fibrinogen binding by *Apoh^-/-^* platelets in response to low thrombin concentrations was blocked by the PAR3 antibody (**Figure 6 C, D**). We also examined activation of WT and *Apoh^-/-^* platelets by the PAR4-activator peptide, AYPGKF, which activates platelets independently of PAR3. We found no effect of β2GPI deficiency on platelet P-selectin expression (**Figure 7A and D**) or fibrinogen binding (**Figure 7B and E**) induced by AYPGKF. Finally, we did not observe any significant difference in ADP-induced activation of *Apoh^-/-^* platelets compared to WT (**Figure 7C and F**), indicating that the effect of β2GPI on platelet activation is specific to thrombin.

**Figure 6.**
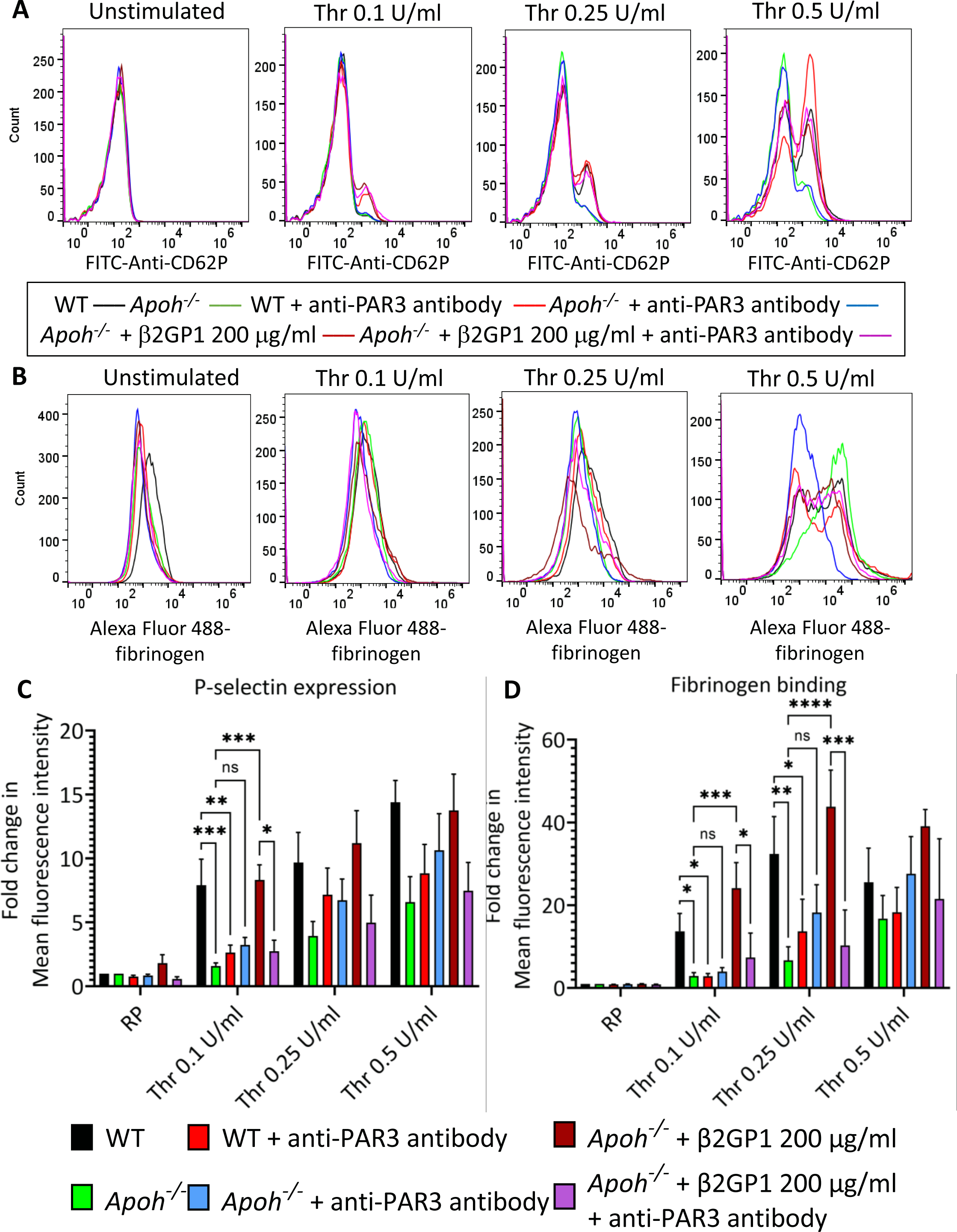
Effect of a PAR3 antibody on thrombin-induced activation of WT and *Apoh^-/-^* platelets. Histogram overlays showing thrombin-induced (A) P-selectin expression and (B) fibrinogen binding of WT platelets, *Apoh^-^*^/-^ platelets and *Apoh^-^*^/-^ platelets supplemented with β2GPI (200 µg/ml) in the presence or absence of an anti-PAR3 antibody (10 µg/ml). (C) and (D) Corresponding bar diagrams for mean P-selectin expression and fibrinogen binding, respectively.

**Figure 7.**
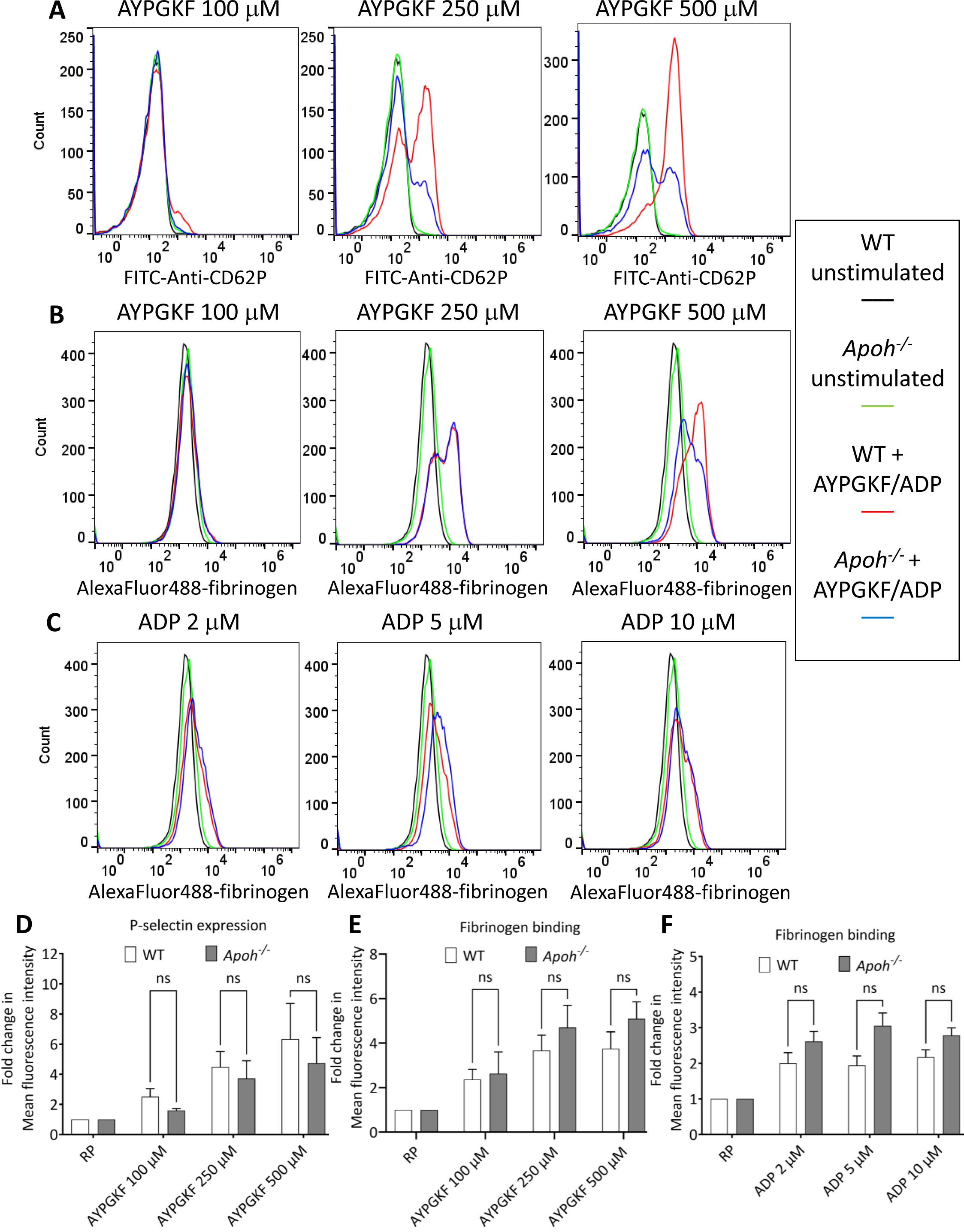
Activation of WT and *Apoh^-/-^* platelets by AYPGKF and ADP. Histogram overlays showing AYPGKF (100-500 µM)-induced (**A**) P-selectin expression and (**B**) fibrinogen binding of WT and *Apoh^-/-^* platelets. (**C**) Histogram overlays showing ADP (2-10 µM)-induced fibrinogen binding of WT and *Apoh^-/-^*platelets. Corresponding bar diagrams for mean (**D**) P-selectin expression and (**E**) and (**F**) fibrinogen binding in response to AYPGKF and ADP, respectively.

Thrombin-induced GPIb signaling is known to synergize with PAR-dependent signaling in mediating platelet activation in response to low-dose thrombin.^43^ However, we did not find any significant inhibition of thrombin-induced P-selectin expression (**Figure 8A and C**) or fibrinogen binding (**Figure 8B and D**) to WT platelets after pretreatment with either O-sialoglycoprotein endopeptidase (OSGE), which cleaves the extracellular domain of GPIbα from the platelet surface^36^ or fibrinogen-γ-peptide, that prevents thrombin-GPIbα interaction.^44^ In concurrence neither OSGE nor fibrinogen-γ-peptide influenced low dose thrombin-induced P-selectin expression (**Supplementary figure 4A**) or fibrinogen binding (**Supplementary figure 4B**) on *Apoh^-/-^* platelets with or without β2GPI supplementation. These findings are not consistent with a role for β2GPI in promoting thrombin-GPIb signaling.

**Figure 8.**
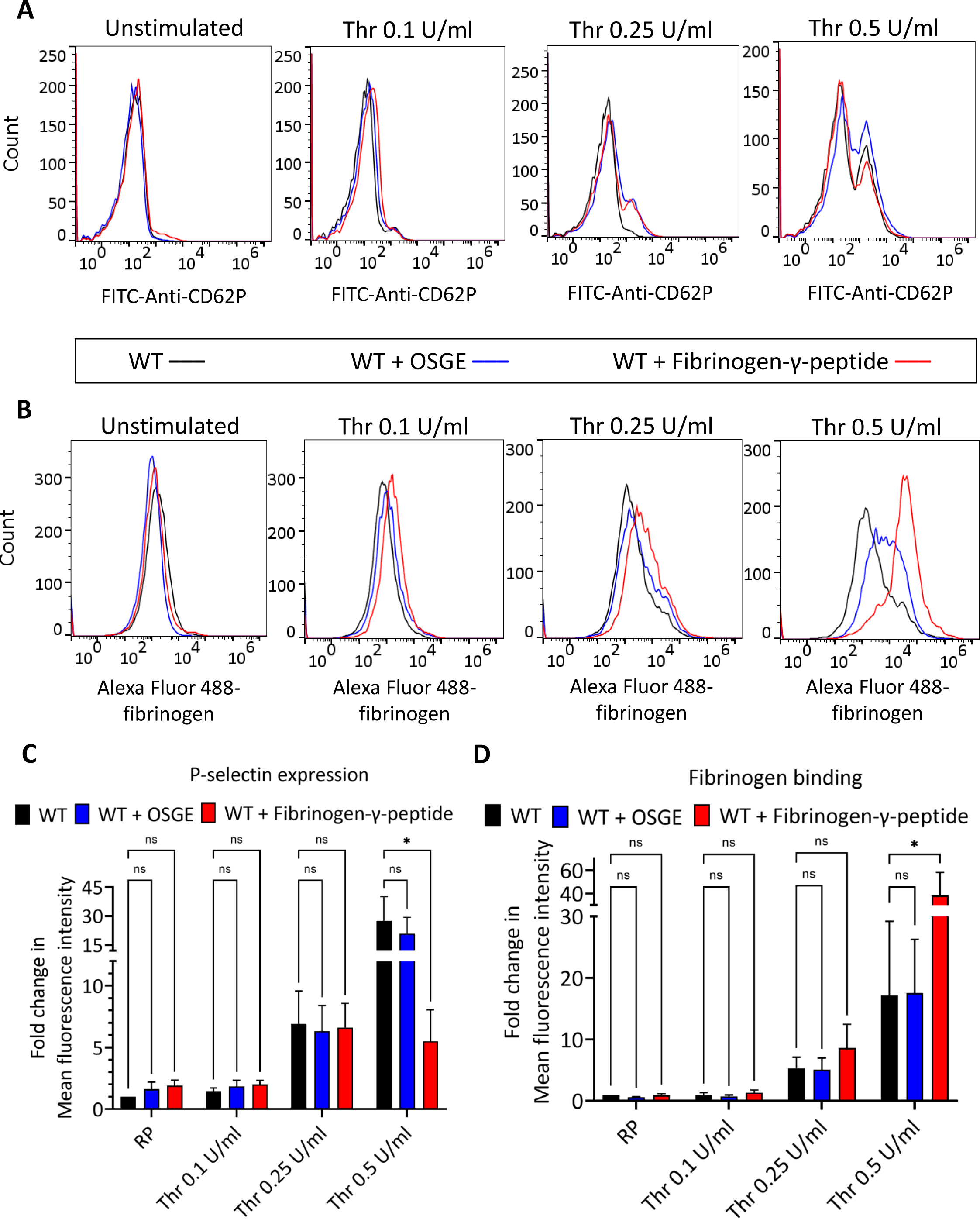
Effect of GP1b signaling inhibition on thrombin-induced activation of WT platelets. Histogram overlays showing thrombin-induced (A) P-selectin expression and (B) fibrinogen binding to WT platelets in the presence or absence of O-sialoglycoprotein endopeptidase (OSGE) (80 µg/ml) or fibrinogen-γ-peptide (200 µM). (C) and (D) Corresponding bar diagrams for mean P-selectin expression and fibrinogen binding, respectively.

## Discussion

β2GPI is an abundant plasma glycoprotein, and a major antigen of APL. Interactions of APL with β2GPI contribute to thrombotic mechanisms in APS^1,2,5,9^, and it is therefore essential to clearly define whether β2GPI itself plays a role in normal hemostasis.

In this study, we used CRISPR/Cas9 to generate *Apoh^-/-^* mice. The deletion generated in *Apoh* was similar to that created by Sheng *et al* using homologous recombination^18^. Although the CRISPR approach does not allow us to target a specific sequence with complete precision^45^, we selected and bred individual progeny of founder mice to generate a clonal population of mice that shared an exon 2-3 deletion in *Apoh* and lacked β2GPI mRNA in the liver and β2GPI in plasma. Our findings demonstrate that mice lacking β2GPI are protected from arterial and venous thrombosis. The consistency of these results using 4 different thrombosis models including an arteriolar intravital imaging model demonstrating early events after vessel injury, WT littermate controls, and extensive sequencing of the deleted locus and adjacent genomic regions in *Apoh^-/-^* mice provides confidence in the fidelity of our models and our experimental results. A major strength of this study is the demonstration that β2GPI promotes thrombosis in the arterial as well as the venous circulation, which is consistent with how patients with acute thrombosis present. Readily observing thrombosis in both the high velocity blood flow environment of arteries and the lower blood flow velocity environment of veins argues against shear stress-mediated thrombosis at the boundary of platelets and endothelial cells.

Despite reports of altered procoagulant activity resulting from interactions of APL with β2GPI, there is relatively little information available concerning the role of β2GPI itself in hemostasis. *In vitro* studies have suggested that β2GPI displays both anticoagulant and procoagulant properties. Reduced levels of β2GPI in patients with DIC has been assumed to reflect consumption, though whether β2GPI may be consumed in the context procoagulant or anticoagulant processes is unknown.^16,46^ β2GPI has been reported to bind thrombin and prolong clotting times in fibrin generation assays by one group^17^, although others found only a modest effect on thrombin generation that occurred following preincubation of β2GPI with lipid at very low tissue factor concentrations.^19^ β2GPI has also been reported to bind FXI and block FXI activation by thrombin or FXIIa^20^, suggesting an anticoagulant effect. Finally, impaired thrombin generation was reported using plasma from β2GPI deficient mice by Sheng *et al*, which is somewhat unexpected since these mice displayed a normal aPTT, dilute Russell’s viper venom time (dRVVT), and dilute kaolin clotting time (dKCT).^18^ Our studies demonstrate that plasma from β2GPI deficient mice supports a normal PT and aPTT. Moreover, there were no differences in thrombin generation induced by either the intrinsic or extrinsic pathways when plasma from WT and *Apoh^-/-^* littermates were directly compared.

Keeling et al reported that β2GPI impaired the activation of protein C in a system comprised of phospholipid vesicles and thrombomodulin.^21^ Additional *in vitro* studies suggested that β2GPI impaired the activated protein C (APC)-mediated inhibition of thrombin generation^47^, and the degradation of FVa.^48^ However, Pozzi *et al* did not find any effect of β2GPI on thrombin-mediated activation of protein C.^17^ Our findings are in agreement with the latter study, as we found no differences in the activation or activity of protein C in plasma from *Apoh^-/-^* mice and WT littermates. The lack of effect of β2GPI on activity of coagulant pathways suggested that other mechanisms, such as an effect on platelet function, might underlie the apparent procoagulant activity of β2GPI. One study reported that binding of β2GPI to washed platelets occurred only in the absence of Ca^2+^; no binding to gel filtered platelets was observed, even after platelet activation by thrombin, collagen, or ADP. In addition, β2GPI inhibited activation of platelets by ADP, but not other agonists.^23,25,26^ Schousboe^24^ reported that incubation of washed platelets with β2GPI decreased adenylate cyclase activity, while Nimpf found that β2GPI increased levels of platelet cGMP in response to ADP and collagen, but not thrombin.^49^ Previous reports have also suggested that β2GPI may impair the prothrombinase activity of resting and lysed platelets, as well as phospholipid vesicles^50^. In many of these studies, however, the platelet activation state following platelet isolation was not well characterized. Moreover, the prolonged tail vein bleeding times that we observed in *Apoh^-/-^*mice suggest that *in vivo*, β2GPI may enhance platelet function.

Our studies demonstrate that platelets from *Apoh^-/-^* mice are less susceptible to activation by low concentrations of thrombin, suggesting deficiencies in PAR signaling as a potential underlying mechanism for the reduced thrombus size in these animals. Human platelets express primarily PAR1 and PAR4^51^, with low levels of PAR3^52^; signaling at low thrombin concentrations occurs through PAR1^51,53^, while at higher thrombin concentrations PAR4 signals synergistically with the P2Y12 receptor.^54^ In contrast to human platelets, mouse platelets express PAR3 and PAR4, but not PAR1^55^, and signaling occurs through PAR4. In isolation, PAR4 signals in response to high concentrations of thrombin, while PAR3 does not signal. However, PAR3 binds thrombin with high affinity through a hirudin-like domain similar to that in human PAR1, and in combination, PAR3 and PAR4 signal at low thrombin concentrations.^55^ This is believed to reflect the ability of PAR3 to bind thrombin at low concentrations, and present bound thrombin to PAR4, leading to PAR4 activation-an example of cofactor (PAR3)-assisted PAR cleavage and signaling.^55^ Our findings suggest that an underlying reason why *Apoh^-/-^* murine platelets fail to activate as well as WT platelets in the presence of low thrombin concentrations may reflect a defect in this system, possibly through impaired thrombin binding to PAR3, impaired interactions between PAR3 and PAR4, or direct PAR4 activation, although normal responses to higher concentrations of thrombin, which directly activate PAR4^55^, as well as AYPGKF, suggest normal PAR4 function (**Figure 5C**).

In support of this hypothesis, we found that a PAR3 antibody that blocks binding of thrombin to PAR3 reduced low-dose thrombin-induced activation of WT platelets, but not *Apoh^-/-^* platelets suggesting that β2GPI is essential for the influence of PAR3 on platelet activation by low thrombin concentrations; this same antibody also blocked the ability of β2GPI to restore the ability of low thrombin concentrations to activate *Apoh-/-* platelets. Moreover, there was neither any effect of β2GPI deficiency on platelet activation by a PAR4-activator peptide, AYPGKF, nor was there any significant difference in ADP-induced activation of *Apoh^-/-^* platelets compared to WT, indicating that the effect of β2GPI on platelet activation is specific to PAR3-facilitated signaling by low-dose thrombin. Other potential explanations are also possible, and additional studies involving analysis of intracellular signaling responses, PAR3 and 4 expression, and potential binding sites of β2GPI on platelets will be required to better define these effects. Thrombin-signaling through GPIb is also known to contribute to platelet activation in response to low dose thrombin.^43^ However, we found that preventing GPIb signaling does not significantly inhibit low dose thrombin induced activation of either WT or *Apoh^-/-^* platelets irrespective of the presence of β2GPI, which indicates that β2GPI is unlikely to promote thrombin-GPIb signaling.

A limitation of this manuscript is that activation mechanisms in murine platelets may differ from those in human platelets. For example, Pozzi et al reported that β2GPI inhibited thrombin-induced aggregation of human platelets by impairing the cleavage of PAR1^17^. Our attempts to study human platelets were hindered by the presence of large amounts of platelet-associated β2GPI, which could not be significantly reduced without multiple washes, making it difficult to isolate β2GPI-depleted platelets without inducing spontaneous activation. In addition, a recent study also demonstrated functional differences in platelets from WT mice compared to those from murine PAR4-deficient mice that expressed human PAR4 on their platelets, suggesting that mouse and human PAR4 may confer differential reactivity to AYPGKF.^56^

Human β2GPI deficiency is rare and does not appear to be associated with a significant bleeding or thrombotic phenotype. In a large study of 812 individuals screened for β2GPI deficiency by Yasuda *et. al*, only two Japanese individuals with homozygous deficiency of β2GPI were detected. These subjects were healthy, though their full medical histories were not provided^57^. Of the 130 Caucasians enrolled in this study, no cases of heterozygous or homozygous deficiency of β2GPI occurred. Likewise, Takeuchi described two siblings, ages 34 and 36, with homozygous β2GPI deficiency but no history of bleeding or thrombotic events.^58^ Though mild prolongation of the Russell’s viper venom time was noted, a comprehensive panel of markers of thrombin generation and fibrin turnover was normal. In another study, Bancsi et al compared the incidence of β2GPI deficiency (defined as β2GPI levels <77% or normal) in a cohort of healthy individuals and subjects with familial thrombophilia^59^. There was no significant difference in the incidence of β2GPI deficiency between these cohorts. There was also no difference in the incidence of low β2GPI levels in patients with protein C deficiency, or between symptomatic or asymptomatic protein C-deficient patients.

In a recent case report, it was speculated that a mutation in β2GPI associated with a mild reduction in plasma β2GPI might account for thrombophilia. However, the proband’s father had the same mutation but no thrombosis, and the proband also had a persistently positive lupus anticoagulant.^60^. Finally, in a large study from China, four *APOH* polymorphisms (c.−32C>A (rs8178822), c.422T>C (rs52797880), c.461G>A (rs8178847) and c.1004G>C) were found to be in linkage disequilibrium.^61^ Further genotyping of 60 individuals from a thrombosis and no thrombosis group demonstrated a slightly higher incidence of the 461G>A variant in the former, leading to an odds ratio of 1.55 for the association of this allele with thrombosis. Mild reduction of β2GPI levels were noted in subjects with this haplotype. However, this report requires confirmation.

A recent study by Passam using β2GPI deficient mice prepared in 2001^18^ reported that these mice displayed a procoagulant phenotype.^30^ However, in contrast to our study, this report did not use littermate controls, and obtained WT control mice from a commercial vivarium different from that in which *Apoh^-/-^* mice were bred. We believe that the well-recognized effects of non-syngeneic mouse genotypes as well as the microbiome on vascular and thrombotic processes might account for the different results between this study and ours.^62–65^

In summary, mice deficient in β2GPI appear to be protected from arterial and venous thrombosis and exhibit a defect in primary hemostasis, suggesting that β2GPI does not function as a natural anticoagulant, at least in mice. In contrast, our studies suggest that β2GPI deficiency is associated with mildly impaired platelet function, which may reflect ability of β2GPI to enhance PAR3-PAR4 interactions in the presence of low thrombin concentrations. The implications of these findings are consistent with reports that patients with β2GPI deficiency have normal hemostasis without evidence of thrombophilia. While it is uncertain whether our studies are directly translatable to human platelet function, we believe they provide important information relevant to murine models of antiphospholipid antibody-mediated thrombosis, as they suggest that the hypothesis that aPL impair natural anticoagulant properties of β2GPI in mice may not be accurate.

## Acknowledgements

This work was supported by HL143402 and HL164516 to KRM, CA223301, AI130131, and HL144113 to AHS, and HL158801 to SJC.

## Authorship and conflict of interest statement

PPK and RKA wrote the initial manuscript draft. RKA generated and characterized *Apoh^-/-^* mice. PPK, RKA and MP performed tail bleeding studies. PPK and MG performed platelet activation studies. GLF and AM performed in vitro coagulation studies and rose bengal and FeCl_3_ arterial thrombosis studies. MH designed and analyzed the cremasteric arteriole thrombosis assays. AAM performed thrombin generation assays. AV and YJS performed the IVC occlusion assay. SS and YJS reviewed the manuscript. AHS conceived and supervised *in vitro* coagulation studies and mouse arterial thrombosis studies and edited the manuscript. SJC supervised platelet activation studies and edited the manuscript. KRM conceived the project, supervised generation of *Apoh^-/-^* mice, and wrote and edited the manuscript.

None of the authors report any conflict of interest with this work.

